# Bridging parametric and nonparametric measures of species interactions unveils new insights of non-equilibrium dynamics

**DOI:** 10.1101/2020.03.02.973040

**Authors:** Chuliang Song, Serguei Saavedra

**Affiliations:** Department of Civil and Environmental Engineering, MIT, 77 Massachusetts Av., 02139 Cambridge, MA, USA; Department of Biology, McGill University, 1205 Dr. Penfield Avenue, Montreal, H3A 1B1 Canada; Department of Ecology and Evolutionary Biology, University of Toronto, 25 Willcocks Street, Toronto, Ontario M5S 3B2 Canada

**Keywords:** Higher-order Interactions, Interaction Matrix, Jacobian Matrix, Model-driven, Model-free, Non-equilibrium Dynamics

## Abstract

A central theme in ecological research is to understand how species interactions contribute to community dynamics. Species interactions are the basis of parametric (model-driven) and nonparametric (model-free) approaches in theoretical and empirical work. However, despite their different interpretations across these approaches, these measures have occasionally been used interchangeably, limiting our opportunity to use their differences to gain new insights about ecological systems. Here, we revisit two of the most used measures across these approaches: species interactions measured as *constant direct* effects (typically used in parametric approaches) and *local aggregated* effects (typically used in nonparametric approaches). We show two fundamental properties of species interactions that cannot be revealed without bridging these definitions. First, we show that the local aggregated intraspecific effect summarizes all potential pathways through which one species impacts itself, which are likely to be negative even without any constant direct self-regulation mechanism. This property has implications for the long-held debate on how communities can be stabilized when little evidence of self-regulation has been found among higher-trophic species. Second, we show that a local aggregated interspecific effect between two species is correlated with the constant direct interspecific effect if and only if the population dynamics do not have any higher-order direct effects. This other property provides a rigorous methodology to detect direct higher-order effects in the field and experimental data. Overall, our findings illustrate a practical route to gain further insights about non-equilibrium ecological dynamics and species interactions.

## Introduction

Community ecology are built upon the idea that species interact either directly or indirectly with other species (Abrams, 1987; Thompson, 2005; Morin, 2009; Vellend, 2016). Indeed, a central theme in ecological research is to understand how species interactions contribute to community dynamics (May, 1972; Pimm, 1982; Allesina and Tang, 2012; Fukami, 2015; Saavedra et al., 2017; Chesson, 2018). Even macro-ecological studies that do not explicitly model species interactions are built upon the idea of an existing balance among species interactions (Hubbell, 2005; Harte, 2011; Staniczenko et al., 2017). Thus, ever since Odum (Odum and Barrett, 2005), most ecologists classify species interactions not by their mechanisms, but according to the effects produced on the growth rate of populations (see Abrams 1987 for an extended discussion on this topic). Yet, this simple definition has different measures and interpretations across theoretical and empirical studies (Case, 2000), making necessary to understand how and when these measures can be linked.

In empirical and theoretical research, the effect of species interactions has been measured following parametric (model-driven) and nonparametric (model-free) approaches (Sugihara, 1994; Turchin, 2003). While the parametric approach has been the cornerstone of quantitative ecology (Kingsland, 2015), the nonparametric approach has been increasingly adopted in empirical studies (Deyle et al., 2016; Ushio et al., 2018; Cenci and Saavedra, 2019; Yu et al., 2020; Bray and Wang, 2020; Karakoç et al., 2020; Ushio, 2020). To explain the differences between the two approaches, we define them using the most general population dynamics of *S* interacting species in the form of continuous ordinary differential equations (the case for discrete difference equations is similar, see Case 2000),

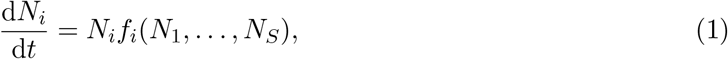

where *N_i_* is the abundance (or biomass) of species *i*, and *f_i_* is the per capita growth rate of species *i*.

The parametric approach typically measures species interactions as *constant direct* effects (mechanistic or phenomenological) between species (Case, 2000; Song et al., 2020), and completely relies on knowledge about the governing population dynamics. The general formalism in the parametric approach partitions the general population dynamics (Eqn. 1) as

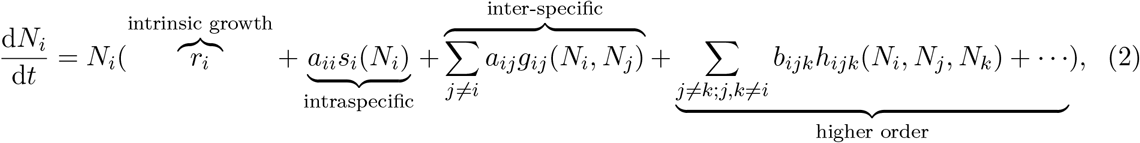

with both pairwise and higher-order terms. The pairwise formalism of population dynamics has been the basis of this approach. The pairwise formalism ignores the higher-order terms and focus only on the pairwise terms, where *r_i_* represents the intrinsic growth rate of species *i* (no density dependency), *a_ij_* represents the constant, direct, intraspecific (if *i* = *j*) and interspecific effect (if *i* ≠ *j*), while *s_i_*(*N_i_*) and *g_ij_*(*N_i_, N_j_*) represent the functional form of the intraspecific and interspecific direct effects, respectively. These constant direct effects *a_ij_* can be the result of indirect mechanisms depending on the level of resolution of the model (MacArthur and Levins, 1967; Abrams, 1987), while the functional forms *g_ij_*(*N_i_, N_j_*) are not restricted to be linear and can incorporate non-additive effects (Billick and Case, 1994; Letten and Stouffer, 2019; Tilman, 1982). A classic example of the pairwise formalism is the Lotka-Volterra (LV) dynamics (Lotka, 1926; Volterra, 1926), where *s_i_*(*N_i_*) = *N_i_*, *g_ij_*(*N_i_, N_j_*) = *N_j_*. The matrix **A** = {*a_ij_*} is called the *interaction matrix*, encoding the strength of pairwise, constant, direct effects (note these effects can be non-additive, see Billick and Case 1994). Regardless of which form of functional responses is used, the sign pattern of the interaction matrix **A** is usually fixed and interpreted as the type of pairwise direct effect, such as: mutualism, competition, predation, or null (Abrams, 1987; Callaway et al., 2002; Chamberlain et al., 2014; Song et al., 2020).

Despite the popularity of the pairwise formalism, the parametric approach can also be applied to a higher-order formalism of the general population dynamics (Eqn. 2; Billick and Case 1994; Kleinhesselink et al. 2019). Higher-order effects correspond to constant direct effects among more than two species (which is fundamentally different from other definitions such as indirect effects or non-additive effects, see Billick and Case 1994). For example, focusing on the higher-order terms in Eqn. 2, *b_ijk_* represents the constant, direct, triple-wise effect. Similarly, *h_ijk_*(*N_i_, N_j_, N_k_*) represents the functional form of the triple-wise direct effect among species *i*, *j*, and *k*—representing the constant change in the per capita growth rate of species *i* under a small change in density of species *j* and *k* (O’Dwyer, 2018; Letten and Stouffer, 2019). Other higher-order direct effects (such as quadruple-wise effect) can be similarly defined (Bairey et al., 2016). Note that the parametric approach, regardless of the specific formalism, can be applied under the assumptions of equilibrium and non-equilibrium dynamics (Case, 2000).

In turn, the nonparametric approach typically measures species interactions as the *local* (state-dependent) *aggregated* (direct and higher-order) effects between two species. Different from the parametric approach, the nonparametric one does not assume any particular governing population dynamics (Sugihara and May, 1990; Ye et al., 2015). Because the local aggregated effect counts all the pathways (including direct and higher-order effects) at a given point in time, it can only be defined pairwise (Deyle et al., 2016; Ushio et al., 2018; Cenci and Saavedra, 2018b). That is, the nonparametric pairwise interaction between two species is measured as the change in the growth rate of species *i* under a small change in density of species *j*. Formally, this can be written as

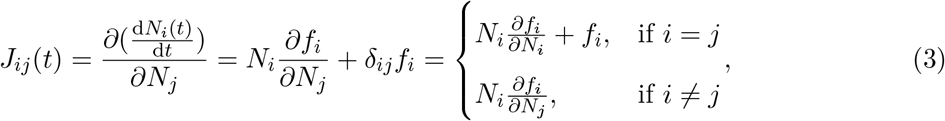

where the Kronecker delta *δ_ij_* is 1 if *i* = *j*, 0 otherwise. The matrix **J** = {*J_ij_*} is called the *Jacobian matrix*. Similarly, this approach can be applied to both equilibrium (May, 1972; Allesina and Tang, 2012) and non-equilibrium dynamics (Sugihara, 1994; Ushio et al., 2018; Cenci and Saavedra, 2019). Note that under equilibrium dynamics, the Jacobian matrix has also been called the *community matrix* (Levins, 1968; May, 1972; Case, 2000; Novak et al., 2016).

Both measures have their own strengths and weaknesses: within parametric approaches, measures have a mechanistic or phenomenological interpretation of a direct effect between species, but the magnitude and dimension of such parametric measures are model-dependent (Cenci and Saavedra, 2018a; AlAdwani and Saavedra, 2019; Letten and Stouffer, 2019). Instead, within the nonparametric approach, measures can be estimated directly from data (such as time series of species abundance) with statistical methods, but they are often hard to be biologically interpreted (Sugihara and May, 1990; Ushio et al., 2018; Cenci et al., 2019). Additionally, regardless of the specific methods, the two approaches hold different conceptualizations about how species interactions can be decomposed: within the parametric approach, measures can be decomposed into intraspecific (the effect of a species on itself), interspecific (the effect of a species on another), and higher-order interactions (the effect of two or more species on another). In contrast, within the nonparametric approach, measures can only be decomposed into intraspecific and interspecific interactions (Deyle et al., 2016; Ushio et al., 2018; Cenci and Saavedra, 2018b). Yet, it remains unclear under what conditions parametric and nonparametric views of species interactions tell a similar story, and what can be learned when they do not coincide.

Importantly, even in equilibrium dynamics, the subtle but central differences in the measure of species interactions between these two approaches have sometimes been a cause of confusion in the literature (Lawlor, 1980; Abrams, 1981). Take the complexity-stability debate as an example, one of the most controversial topics in theoretical and community ecology (May, 1972; McCann, 2000; Ives and Carpenter, 2007; Landi et al., 2018; Xu et al., 2019). As it has been shown (Logofet, 2005), much of the debate has been generated by aiming to generalize ecological dynamics and species interactions under a nonparametric approach. However, the merger between parametric and nonparametric approaches to species interactions in such a context is only possible under the (often implicitly) assumption of a LV model and equal equilibrium states for all species (Haydon, 1994; Novak et al., 2016; Vázquez et al., 2007). While researchers have been increasingly recognizing these assumptions in equilibrium dynamics (Berlow et al., 2004; Novak et al., 2016), it remains unclear whether the two approaches can be transferable in non-equilibrium dynamics, and more importantly, whether the transferability may reveal hidden ecological dynamics.

Here, we revisit and show how to bridge two of the most used measures of species interactions across the parametric and nonparametric approaches. We show that bridging parametric and nonparametric approaches present new ecological insights that cannot be revealed without this bridging. Specifically, we study species interactions under three categories: intraspecific, interspecific, and higher-order interactions. In the reminder, we begin by showing that the measures in parametric and nonparametric approaches can be linked if and only if all species interactions are pairwise (i.e., no higher-order interactions present) regardless of the dynamics assumed. Next, we demonstrate that interspecific interactions are more transferable across measures than intraspecific interactions. Next, we show two applications by building on the differences between approaches. Finally, we discuss how and when these measures can be combined to gain further insights about non-equilibrium ecological dynamics and higher-order interactions.

## The translucent mirror between measures

### Intraspecific interactions

Under the parametric approach, a negative, constant, direct, intraspecific effect *a_ii_* is often considered as *self-regulation* or *intraspecific density dependence* (Case, 2000). However, under the nonparametric approach, the interpretation of the local aggregated intraspecific term *J_ii_* is more complicated. For example, following the general parametric formalism defined in Eqn. (2), the elements of the Jacobian matrix are defined as

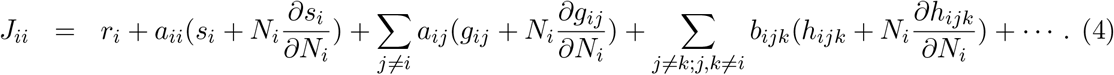

and when the system is at equilibrium, it reduces to

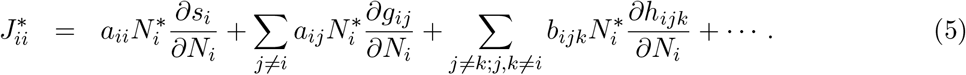

Note that the notation for *s_i_*, *g_ik_* and *h_ijk_* in Eqn. 4 and thereafter has been simplified, but they are still functions of the species abundances *N*. Therefore, regardless of the presence of higher-order effects (whether *b_ijk_* are all zeros) or the system is at the equilibrium 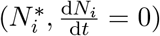, the term *J_ii_* measures the local aggregated effect across all the pathways under which species *i* can affect itself (not only the direct self-loop from *i* to *i*).

Hence, it is natural to ask what is the link between *a_ii_* and *J_ii_*. In general, a negative sign in *J_ii_* does not imply a constant direct self-regulation (*a_ii_* < 0), and vice versa (Somorjai and Goswami, 1972; Haydon, 1994). This property can be easily illustrated using the logistic population dynamics of a single species,

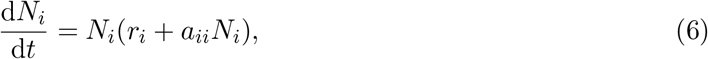

where *r_i_* and *a_ii_* correspond to the intrinsic growth rate and the direct self-regulation of the single species *i*, respectively. At the equilibrium 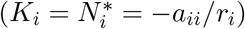, the constant, direct, intraspecific effect is given by *a_ii_*, which is interpreted as a constant self-regulation. In turn, from Eqn. (5) the Jacobian *J_ii_* equals 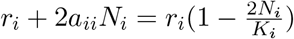, which is always positive when *N_i_ < K_i_/*2, negative otherwise. This implies that, in general, the interpretation of intraspecific interactions across the parametric and nonparametric are not the same.

Then, when can *J_ii_* be transferable into *a_ii_*? If we require that the signs of *a_ii_* and *J_ii_* be the same, we need the system at equilibrium following LV dynamics. The reasoning is that, no higher order effects exist in LV dynamics (i.e. *b_ijk_* = 0) and the partial derivative of *g_ij_* with respect to *N_i_* is 0 (i.e. 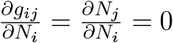), thus that the second and third terms on the right-side of Eqn. (5) vanish. If we additionally require that *a_ii_* = *J_ii_*, then on top of the two previous requirements, we need all equilibrium abundances to be exactly the same (May and Mac Arthur, 1972; Song and Saavedra, 2018). While it is not explicit, note that previous work (May, 1972; Coyte et al., 2015) on the complexity-stability debate operates under these assumptions.

### Interspecific interactions

Assuming that all direct effects are pairwise as described in Eqn. (2), the local, aggregated, interspecific effect can be derived as

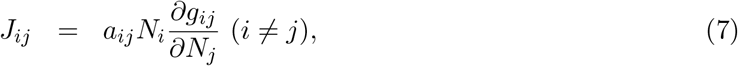

which only includes the direct effect (functional form) *g_ij_* between species *i* and *j*. Under this assumption, *J_ij_* and *a_ij_*(*i* ≠ *j*) always have the same sign because 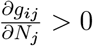 (biologically, this means that effects are stronger with larger species abundances).

Instead, assuming that constant direct interactions include higher-order effects as in Eqn. (2), the Jacobian (the local aggregated effects) can be derived as

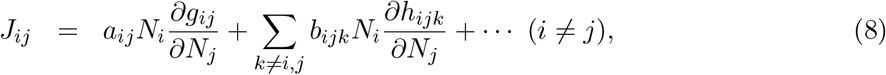

where *J_ij_* encodes not only the direct interspecific effects, but also the higher-order effects coming from species other than species *i* and *j*. Therefore, *J_ij_* can be interpreted as the local (state-dependent) direct effect between species *i* and *j if and only if* all (parametric) direct effects are pairwise. That is, under higher-order effects, there is no simple link between (parametric) *a_ij_* and (nonparametric) *J_ij_* interspecific interactions. This also shows that the interspecific *J_ij_* (*i* ≠ *j*) is fundamentally different from the intraspecific *J_ii_*.

## Learning from the differences between approaches

### Debates on self-regulation and stability

Importantly, the differences between approaches (measures) can offer an opportunity to gain further insights about non-equilibrium ecological dynamics and higher-order interactions without modeling them (AlAdwani and Saavedra, 2019). For example, focusing on dynamics and building from the classic complexity-stability debate (May, 1972), it is assumed that a community can be dynamically stable only if most of the constant, direct, intraspecific terms are negative (*a_ii_* < 0), i.e., if “the population of each species would by itself be stable” (May, 1972). This assumption comes from the observation that dynamical stability requires that most of the local, aggregated, intraspecific terms are also negative (*J_ii_* < 0) (May, 1972; Yodzis, 1980; Sterner et al., 1997; Moore and de Ruiter, 2012; McCann, 2011; Barabás et al., 2017). Yet, there is few empirical evidence to support the addition of direct self-regulation (*a_ii_* < 0) for primary consumers and top predators (Pimm and Lawton, 1977; Tilman, 1982; Chesson, 2013), which would make most systems unstable.

This apparent contradiction arises from the ill perception that a negative *J_ii_* requires a negative *a_ii_*. However, recalling the link between *J_ii_* and *a_ii_* (Eqn. 4), *J_ii_* can be expressed in the absence of self-regulation (*a_ii_* = 0) as

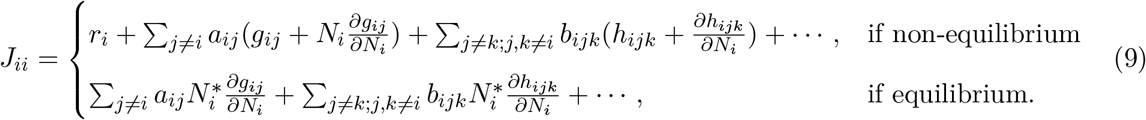

This implies that a negative *J_ii_* (in equilibrium and non-equilibrium dynamics) in a non-self-regulated species *i* (i.e., *a_ii_* ≥ 0) can arise in a broad class of nonlinear ecological dynamics simply by satisfying two conditions (Song et al., 2018): (1) a negative intrinsic growth rate (i.e., *r_i_* < 0), and (2) at least one negative, constant, direct, interspecific effect (i.e., *a_ij_* < 0). Note that those conditions do not apply to LV dynamics in equilibrium because of the linearity of the dynamics (i.e.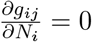). Figure 1 shows a simple example in a 3-species food chain: both the consumer and the top predator have no constant, direct self-regulation; yet they can exhibit negative, local, aggregated, intraspecific effects. In contrast, the primary producer does have constant, direct self-regulation; yet it does not always exhibit a negative, local, aggregated, intraspecific effect. Hence, apart (or instead) of local aggregated self-regulation mechanisms, these (or other conditions) can be taken as stabilization sources of ecological communities.

**Figure 1:**
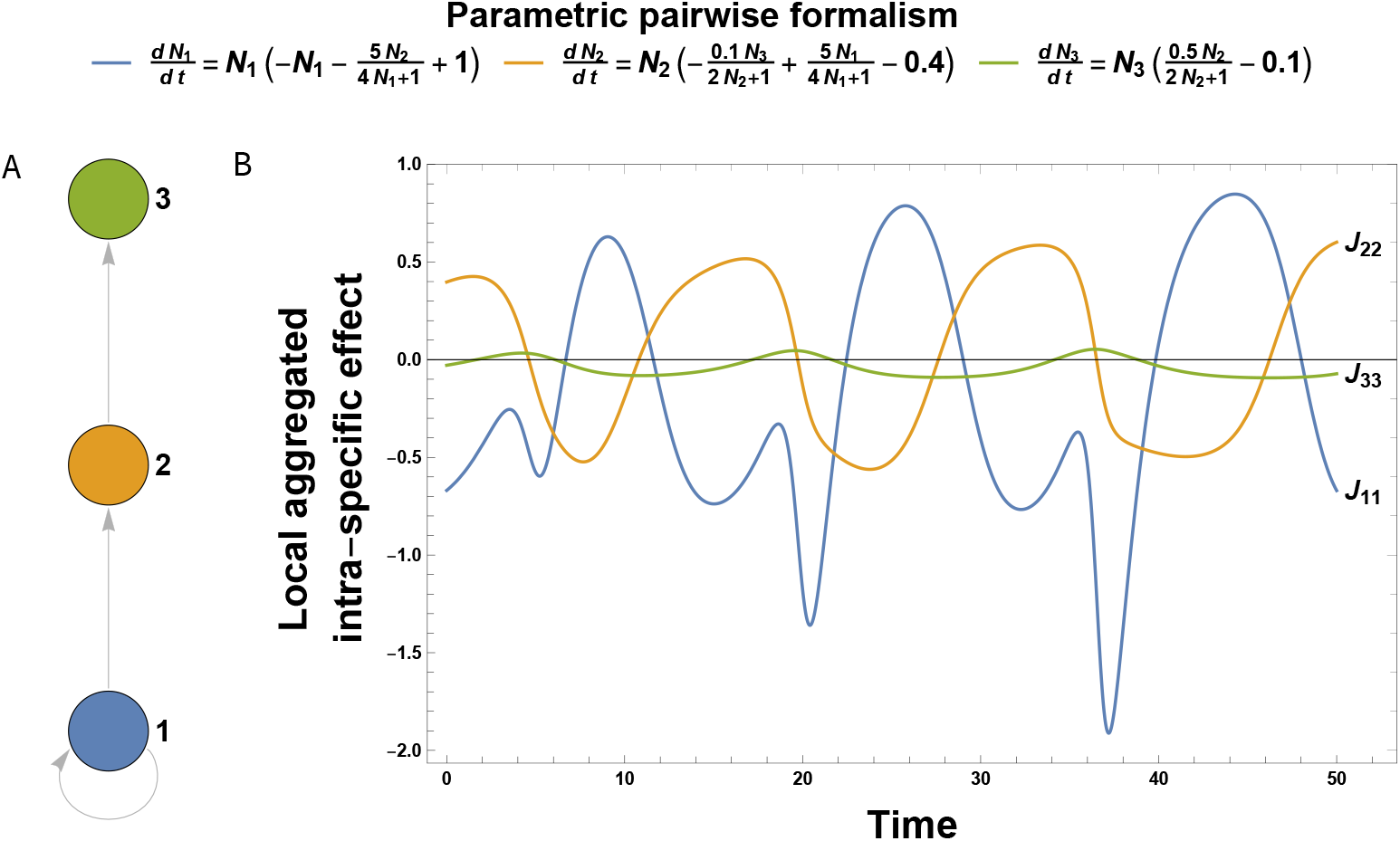
Local, aggregated, intraspecific effects can be negative without a constant, direct, self-regulation mechanism. Panel (A) shows a simple 3-level trophic chain with a primary producer (bottom circle), a consumer (middle circle), and a top predator (top circle). These species are linked by arrows showing the standard energy/biomass flow. Note that only the primary producer has a constant direct self-regulation (typically used in parametric approaches), i.e., *a*_11_ < 0, whereas *a*_22_ = *a*_33_ = 0. The governing equations describing the population dynamics of the 3-species trophic chain are shown on the top. Panel (B) shows the local (state-dependent), aggregated, intraspecific effects *J_ii_* (typically used in nonparametric approaches) when the trophic chain is governed by a type II functional response (parameters are taken from Ref. (Hastings and Powell, 1991)). Top predator (*J*_33_) shows mostly negative, local, aggregated effects to itself; whereas both the consumer (*J*_22_) and the primary producer (*J*_11_) show anti-correlated oscillatory sign patterns of local, aggregated effects to themselves.

### Detection of higher-order interactions

Ecology has seen the re-emergence of interests in higher-order interactions (Mayfield and Stouffer, 2017; Grilli et al., 2017; Levine et al., 2017). However, it remains challenging to convincingly detect the presence of higher-order interactions in empirical data (Kleinhesselink et al., 2019; Letten and Stouffer, 2019; Xiao et al., 2020). The different interpretations of *J_ij_* in the presence of higher-order effects provide a new method to detect their existence. For example, it has been found that *J_ij_* can change its sign across time in a community (Ushio et al., 2018). If we assume that the governing population dynamics only consists of pairwise direct effects (Eqn. 2), then this result should be interpreted as the change of the type of the constant, direct, interspecific effect (i.e., the sign of the parameters in the governing population dynamics have to change). However, if we assume that the governing population dynamics is fixed, then this result should be interpreted as the presence of higher-order direct effects. Figure 2 shows a simple 3-species competing system with and without higher-order direct effects that illustrates these points. Of course, the assumption relies on our belief of how nature operates. For example, previous work (Ushio et al., 2018) has assumed that the governing population dynamics is fixed, implying the presence of higher-order direct effects.

**Figure 2:**
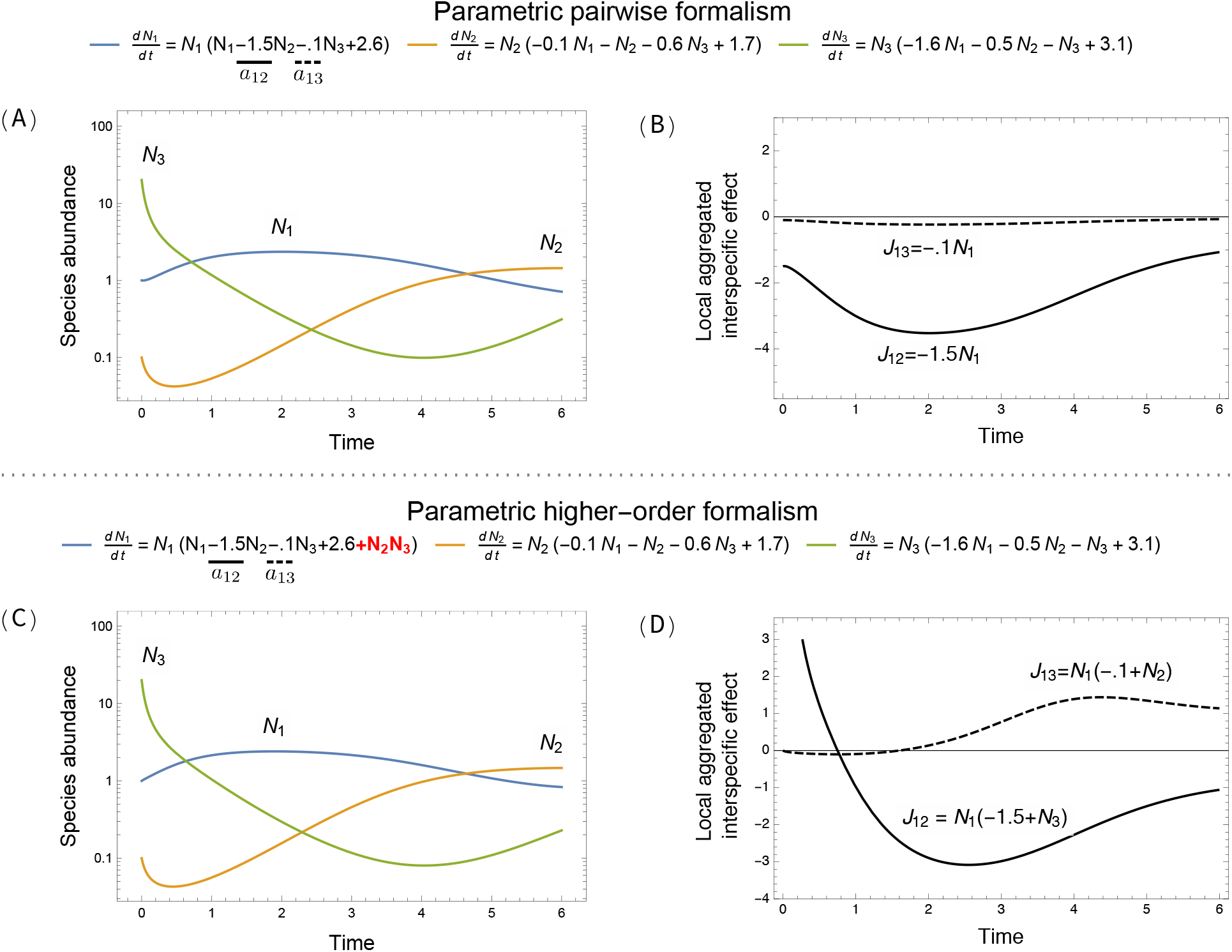
Local, aggregated, interspecific effects and constant, direct effects have the same sign if and only if all direct effects are pairwise based (i.e., absent of higher-order direct effects). Panels (A)-(B): Three competing species are governed by Lotka-Volterra dynamics without any higher-order direct effects (shown above panels; parameters adopted from Saavedra et al. 2017). Panel (A) shows the time series of species abundances. Panel (B) shows the corresponding local, aggregated, interspecific effects (typically used in nonparametric approaches) that species 2 and 3 have on species 1, where both *J*_13_ and *J*_12_ are always negative. Panels (C)-(D): Three competing species are governed by Lotka-Volterra dynamics with an added higher-order direct effect from species 2 and 3 on species 1 (highlighted in red; shown above panels). This formulation of higher-order effects is conceptually inspired by Levine et al. 2017 and mathematically adopted from Letten and Stouffer 2019). Panel (C) shows the time series of species abundances, which exhibit similar patterns as the model without higher-order interactions shown in Panel (A). However, Panel (D) shows that both *J*_12_ and *J*_13_ change their sign, which are fundamentally different from the patterns shown in Panel (B).

## Discussion

Traditionally, the parametric and nonparametric approaches have considered different measures and interpretations of species interactions. That is, species interactions are measured as constant-direct and local-aggregated effects within the parametric and nonparametric approaches, respectively. However, their interpretations have been occasionally used interchangeably (e.g., when describing the stability conditions of an ecological community (May, 1972; Coyte et al., 2015)), limiting our opportunity to use their differences to gain new insights about ecological systems. In this line, here we have provided a bridge between these two approaches (measures) and illustrated its utility. In particular, we have shown three fundamental properties of species interactions. First, the local, aggregated, intraspecific effect summarizes all potential pathways through which one species impacts itself, which can be negative without any direct self-regulation mechanism (see Fig. 1). Second, the local, aggregated, interspecific effect only measures the direct effect between two species if and only if the population dynamics does not have any higher-order direct effects (see Fig. 2A-B). Third, higher-order direct effects can be detected by studying changes of interaction signs within a nonparametric approach (Figure 2C-D).

Species interactions are a multidimensional concept (Callaway et al., 2002; Nakazawa, 2020), which naturally resulted in multiple definitions, ranging from mechanistically motivated characterizations to highly phenomenological representations (White and Marshall, 2019). However, despite the fact that these definitions are distinct mathematical entities, their construction implies that they must be inherently linked given that they all describe properties of species interactions. Importantly, most of the definitions can be classified as either parametric or nonparametric. The parametric approach decomposes species interactions in *biologically interpretable* intraspecific, interspecific, and high-order direct effects. In turn, the nonparametric approach decomposes species interactions in *computationally feasible* intraspecific and interspecific aggregated effects. Therefore, instead of linking specific definitions case-by-case, we have bridge these two approaches by focusing on their high-level conceptual links.

We hope this *Forum* paper can open a dialogue between the parametric and the nonparametric approaches. The parametric approach has dominated community ecology (Kingsland, 2015), while the nonparametric approach has recently received increasing attention in the past decade (Deyle et al., 2016; Ushio et al., 2018; Cenci and Saavedra, 2019; Yu et al., 2020; Bray and Wang, 2020; Karakoç et al., 2020; Ushio, 2020). While both approaches have shaped our understanding of ecological dynamics, little is known about when and how we can transfer the knowledge from one approach to the other. Importantly, we have shown that the transferability is necessary and provides a new perspective that each approach itself cannot offer. For example, the Achilles’ heel of the parametric approach is to evaluate whether the model has included enough details of the system under investigation. Indeed, if we assume a pairwise formalism, while the system is actually governed by a high-order formalism (Box 1), then we are likely to make false predictions of the system (Letten and Stouffer, 2019). However, the computational methods emerging from the parametric approach are difficult to distinguish (e.g., functional responses and higher-order interactions) (AlAdwani and Saavedra, 2019). Yet, relying upon the computational feasibility of the nonparametric approach (Martin et al., 2018; Deyle et al., 2016; Cenci and Saavedra, 2019), we may be able to distinguish the nature of species interactions acting on a system. Therefore, we believe that a better understanding of both the measures and assumptions used across parametric and nonparametric approaches can improve our knowledge of species interactions and ecological dynamics in general.

## Competing interests

The authors declare no competing interests.

## Notes

### Competing Interest Statement

The authors have declared no competing interest.

### Summary of Updates

We have reorganized the text to clarify our key contributions.

